# Knowledge, attitude and practice of secondary school students toward COVID-19 epidemic in Italy: a cross selectional study

**DOI:** 10.1101/2020.05.08.084236

**Authors:** Dafni Souli, Maddalena Dilucca

## Abstract

The coronavirus disease (COVID-19) is a highly transmittable and pathogenic viral infection caused by Severe Acute Respiratory Syndrome Coronavirus 2 (SARS-CoV-2), which emerges in December 2019 in Wuhan, China and spreads around the world at the beginning of 2020. The World Health Organization declares the outbreak a Public Health Emergency of International Concern on the 30th of January, and a pandemic on the 11th of March. On the 4th of March, the Italian government orders the full closure of all schools and universities nationwide.

The aim of this study is to investigate the knowledge, practice and attitudes (KAP) of secondary school at the time of COVID-19 pandemic in Italy. In this cross-sectional, web-based survey, conducted among the high school student population with age ranging from 14 to 19 years old, a questionnaire with 19 items regarding the KAP toward COVID-19 is asked. Study participants are recruited from several secondary schools of different areas. Frequencies and histograms are computed for descriptive purposes. Statistical analysis is computed with Chi square test, utilized to depict relevant difference between geender. Among a total of 2380 students who answers the questionnaire, 40.7% are male and 59.3% are female.

Level of knowledge about generical characteristics of COVID-19 is quite similar among gender. Students present a good level of knowledge about the clinical presentation of the disease, the basic hygiene principles, the modes of transmission and the method of protection against virus transmission. The knowledge about number of this pandemia and easy scientific correlation with COVID-19 is quite confused. The most frequently reported source of knowledge about COVID-19 is television, whereas the less is the school. Our findings suggest that student population shows appropriate practice, and positive attitude towards COVID-19 at the time of its outbreak.

More emphasis should be placed on education of the student partecipants about biological meaning of this infection and relative preventive or future measures.

## 1 Introduction

Coronavirus disease 2019 (officially known as SARS-CoV-2 or COVID-19) is an emerging acute respiratory illness that is caused by a novel coronavirus and is first reported in December 2019 in Wuhan, Hubei Province, China, [1] from where it spread rapidly to over 198 countries [2, 3]. In response to this serious situation, the World Health Organization (WHO) declares it a public health emergency of international concern on 30th of January and calls for collaborative efforts of all countries to prevent the rapid spread of COVID-19. It is declared as a global pandemic by WHO on 12th of March [4, 5].

The outbreak of coronavirus in Italy is officially confirmed to be on 31th of January, when two Chinese tourists in Rome tested positive for the virus [6]. One week later an Italian man repatriated back to Italy from the city of Wuhan, China, is hospitalised and confirmed as the third case in Italy [7]. A cluster of cases is later detected, starting with 16 confirmed cases in Lombardy [8]. As of 7th of May 2020, over three million cases of COVID-19 has been reported with a death toll of over 270.000 patients [9]. Among the top ranking countries, Italy is reported to be in the third position, after America and Spain, with over 215.000 confirmed cases and over 26.000 deaths [9].

Health authorities in Italy have made substantial efforts to control the disease through various measures. On 8th of March, Prime Minister of Italy extends the quarantine lockdown to cover the whole region of Lombardy and 14 other northern provinces. On 10th of March, he increases the quarantine lockdown to cover all of Italy, including travel restrictions and a ban on public gatherings [10]. Public education is considered as one of the most important measures that could help control the diseases, as has been the case regarding MERS [11] or SARS [12, 13]. In fact, On 4th of March, the government announces the closure of all degree of schools.

The main aim of our present study is to investigate the level of knowledge, attitude, and practice of young subjects attending secondary schools toward COVID-19 infection. This study is interesting to detect variables associated with a satisfactory level of them and to explore how knowledge about the disease has affected their lifestyles or health behaviors.

## 2 Materials and Methods

### 2.1 Questionnaire preparation

For the purpose of this study, a questionnaire self-administered is developed especially for this research in native spoken language (Italian). A small subgroup of twenty male and twenty female students are asked to complete the questionnaire and then to ask some questions on whether the questionnaire is easy to understand, to complete and to submit. Reliability of the questionnaire in its translated form is measured by calculating Cronbach’s alpha for each total scale. In our case, a value of Cronbach’s alpha > 0.8 is considered significative.

After validation questionnaire, we develop the final version with 19 items, featured in the form of a multiple choice answers. The questionnaire consists of two parts: demographics and KAP [14]. The questionnaire is subdivided as following questions: demographic information (items 1-4), knowledge of signs and symptoms of the disease, the methods of transmission of the disease and of prevention (items 5-16), the impact of the disease orientation on participants’ lifestyles and their sources of information (item 17-19) [15]. This takes approximately 5 minutes to fill out.

### 2.2 Data collection

This web-based survey is carried out through various social media platforms. Through the link, the participants can view the questions simply by clicking on it and answer. The cover page of the questionnaire includes a short introduction regarding the objectives, the procedures, the voluntary nature of participation, the declarations of confidentiality and anonymity. The inclusion criteria regard Italian nationality younger with age ranging to 14 from 19 years old attending secondary high school.

The questionnaire is answered by over 2300 participants anonymously from the 1st to the 5th of May, in the first days of the second phase in Italy (it begins on the 4th of May). Demographic variables are recorded along with other factors regarding the populations’ knowledge, attitude, practice, and risk assessment concerning COVID-19 [16].

### 2.3 Statistical analysis

All the statistical analyses are performed by using statistical package for social sciences (SPSS Inc., Chicago, Illinois, USA) version 26.0. Data are presented as mean ± SD, frequencies of knowledge answers and proportions as appropriate. The Chi-square test is used to compare categorical data between gender. The statistical significance level is considered with p-values < 0.05 (two-sided) [17].

## 3 Results and discussion

### 3.1 Demographic information

Out of 2380 students who answer the questionnaires, 1410/2380 (59.3%) are female and 970/2380 (40.7%) are male. Students attend high school from five areas of Italy: Lombardia (Milan), Lazio (Rome and Frosinone), Puglia (Lecce), Campania (Naples) and Calabria (Catanzaro). The age of participants ranges from 14 to 19 years old, with a mean age equal to 17 and SD 3.64 (for more details, Figure 1 shows the demographic characteristics of the participants). 770/2380 (32.3%) of the participants attends scientific school, 90/2380 (3.8%) the artistc, 590/2380 (24.8%) the classical, 20/2380 (0.8%) human science, 530/2380 (22.3%) linguistic and 130/2380 (5.4%) the techincal one (see Figure 2).

**Figure 1.**
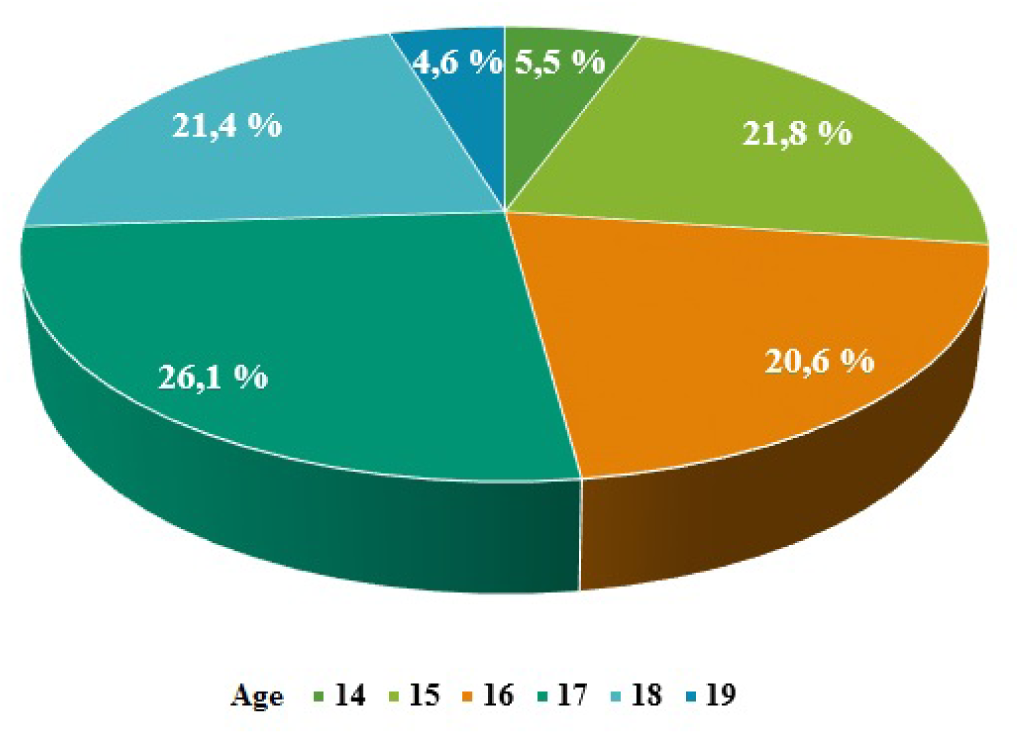
Distribution of age in dataset. The age of participants ranges from 14 to 19 years old, with a mean age equal to 17 and SD 3.64.

**Figure 2.**
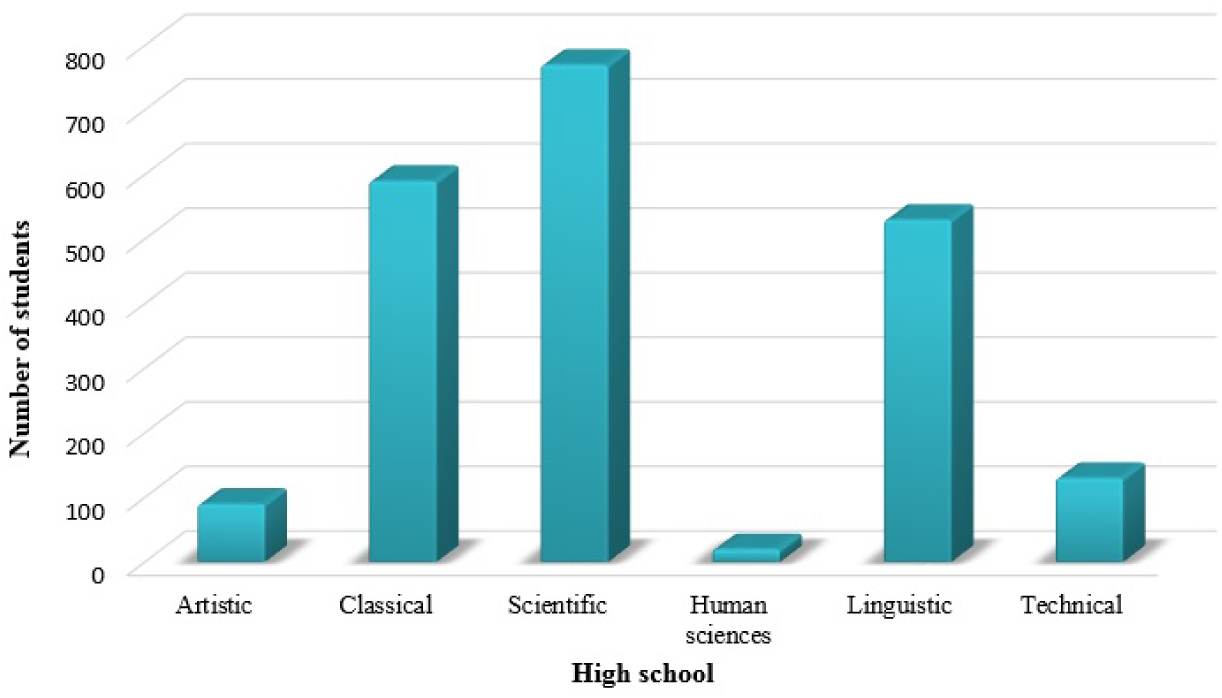
Histograms of students in High school. 770/2380 (32.3%) of the participants attends scientific school, 90/2380 (3.8%) the artistc, 590/2380 (24.8%) the classical, 20/2380 (0.8%) human science, 530/2380 (22.3%) linguistic and 130/2380 (5.4%) the techincal one.

### 3.2 Kknowledge of COVID-19

Basic information about COVID-19 are required to participants. To the first question ”What is COVID-19?” 1690/2380 (71%) correct answer that it is a virus, 523/2380 (22%) respond that is a classical flu, whereas only 121/2380 (5%) respond that is a bacteria. The remaining does not respond [18]. The wrong information on the difference about COVID-19 between bacterium and virus is probably due to the fact that not all students have studied these differences in subject Science, either for their age (class attended) or for their kind of study. As regards the confusion with the normal flu this may be due to the initial media advertisement with which the COVID-19 was introduced.

Knowledge about clinical presentation of the COVID-19 is detailed in 2. Among students of both genders, there is an overall agreement about responses. However, 90.7% of female students respond against 76.2 % of male students. Female show a slightly better understanding of clinical presentation of COVID-19 (see percentage of answers of fever in the following Table 1). It is strange to note that more than a quarter rispectevely of males and females links the phenomenon of COVID-19 to nasal congestion. This wrong answer may be due to the common use of linking fever with symptomes of a cold [19, 20].

**Table 1.**
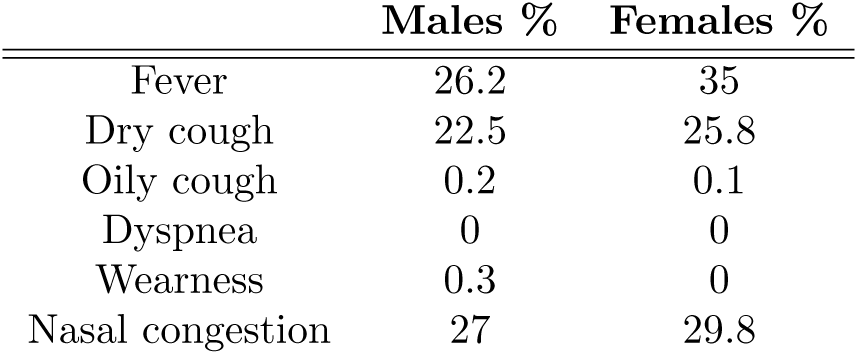
Knowledge about clinical presentation of the COVID-19. Chi-square test is calculated to measure differences between genders and all distributions well superate the test.

Table 2 details knowledge about modes of transmission of COVID-19. There is a similar level of knowledge about mode of transmission among gender. The most frequently reported source of transmission is exposure via coughing and sneezing. The proportion of students who thinks hand shaking is a mode of transmission is lower than those who think touching surfaces might increase risk of infection. Moreover, only 0.1% of the participants thinks that consuming aliments increases risk of infection and only 0.2% thinks that the contact of animals is dangerous for human. It is interesting to show as 2% of males and 0.3% females thinks that the risk of infection is connected by being outdoors.

**Table 2.** Knowledge about modes of transmission of the COVID-19. Chi-square test is calculated to measure differences between genders and all distributions well superate the test.

|  | Males % | Females % |
| --- | --- | --- |
| By being outdoors | 2 | 0.3 |
| By touching surfaces | 38 | 35 |
| By animals to human | 0.2 | 0.3 |
| Via coughing and sneezing | 45 | 49 |
| By consuming aliment | 0.1 | 0 |
| Via hand shaking | 10 | 12 |

As illustrated in Table 3, the most frequently reported method of protection against virus transmission is hand washing. Nevertheless, a higher proportion of students remarks the importance of wearing tissues protect hands. Almost no percentage believes that it is appropriate to touch the nose or eyes or not wear face mask protection. Even in these questions, there is a similar level of knowledge method of protection against virus transmission among gender [21].

**Table 3.** Knowledge method of protection against virus transmission COVID-19. Chi-square test is calculated to measure differences between genders and all distributions well superate the test.

|  | Males % | Females % |
| --- | --- | --- |
| Hand washing | 58 | 87 |
| Using tissues protect hands | 22 | 12 |
| Not using face mask protect | 0.1 | 0 |
| Touching nose and eyes | 0.5 | 0.3 |

According to recent statistics, COVID-19 affects more the male population. So, we ask students if they know about this information: 66.8 % of them answers correctely, 8% answers that females risk more and the remaining replies that does not know. An other question is about the number of deaths with COVID-19 in world and Italy. We know that estimating the correct number of deaths is a sensitive topic especially related to the number of swabs made on patients and so, we consider correct answers only those relating to official data (see website [9]). Unfortunately, only 15% responds correctly, identifying at least the order of measurement of the number both nationally and worldwide. Then we consider which age group is most affected in the population (see Figure 3). 120/2380 (5%) answer that people with class of age 50-60 risk a lot, 720/2380 (30.2%) people with 60-70, 1040/2380 (43.7%) people 70-80 and 500/2380 (21%) people over 80. We could confront our histogram of Figure 3 with official data to understand that students answer incorrectly in this question. In Italy major percentage of deaths is in class over 80 and the second class is 70-80. 2306/2380 (96.9%) students answer that the area most affected in Italy is Lombardia (correct answer), whereas 74/2380 (3%) answer that is Piemonte.

**Figure 3.**
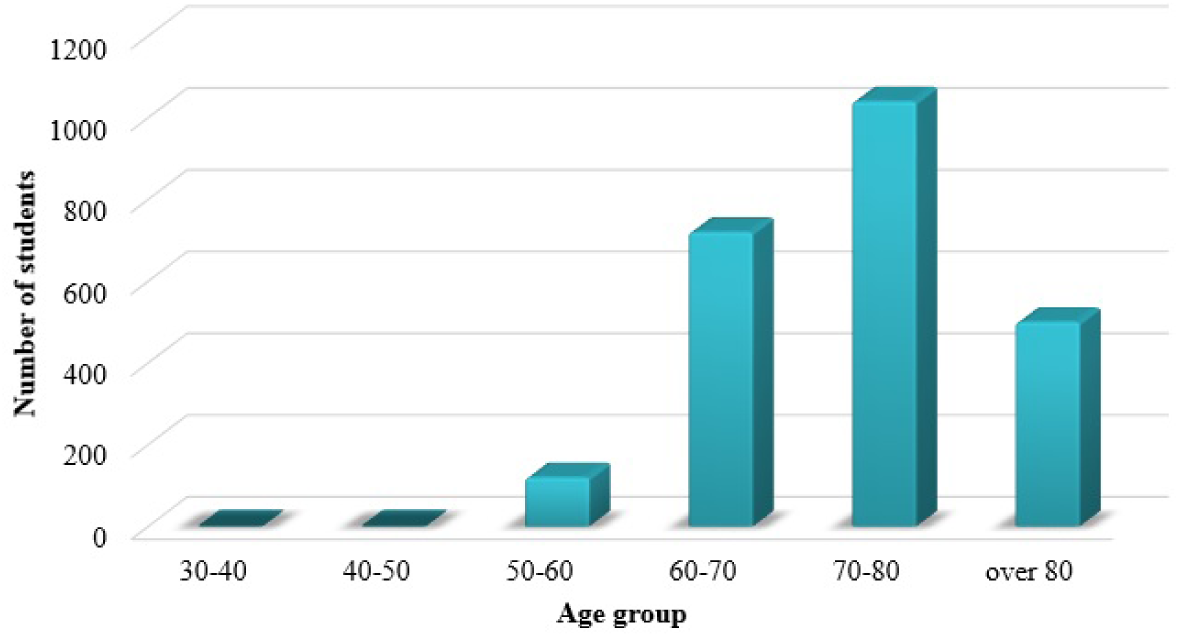
Number of students that answers to questions which age group is most affected in the population.

### 3.3 Attitude towards COVID-19

In the second part of the questionnaire, we ask for generic information about COVID-19 vaccines [22].2058/2380 (86.5%) answer that does not still exist a vaccine, 123/2380 (5.2%) answer that they do not know and the remaining 8.3% that vaccine exists and that soon it will be possible to do it. We know that COVID-19 vaccine is a hypothetical vaccine against coronavirus disease. Although no vaccine has completed clinical trials, there are multiple attempts in progress to develop such a vaccine. In April 2020, 115 vaccine candidates are in development [23, 24]. On 7th of May first COVID-19 vaccine test on animals successful.

2330/2380 (97.9%) claim to know what antibodies are, but in the next question when they have to answer how to check for antibodies, only 1537/2380 (64.6%) answer with blood analysis; the remaining answers pharyngeal swab or urine analysis. 119/2380 (5.1%) answer that flu is a necessary symptome to check antibodies. We remember that antibodies are central to the body’s response to a viral infection. Once they have developed, they can protect the individual from becoming ill after re-infection by a certain pathogen. Knowing which antibodies are developed against COVID-19 helps to understand who has been infected with the virus [25].This type of test is a serological (blood) test and documents the presence of antibodies produced by the immune system against SARS-CoV-2.

### 3.4 Source of information regarding COVID-19

Instead, the source of the individuals’ information about COVID-19 is recorded. It includes social media and internet, news media (TV, magazines, newspapers), family, friends, school and health-care providers, such as doctors. The most frequently reported source of knowledge about COVID-19 is television (see Figure 4). Facebook is the most frequently cited source of knowledge among social media followed by Whatsapp and Instagram. About 20% of the participants reports visiting the website as a source of knowledge. Against other media, school appears to be a less favoured option forgathering knowledge.

**Figure 4.**
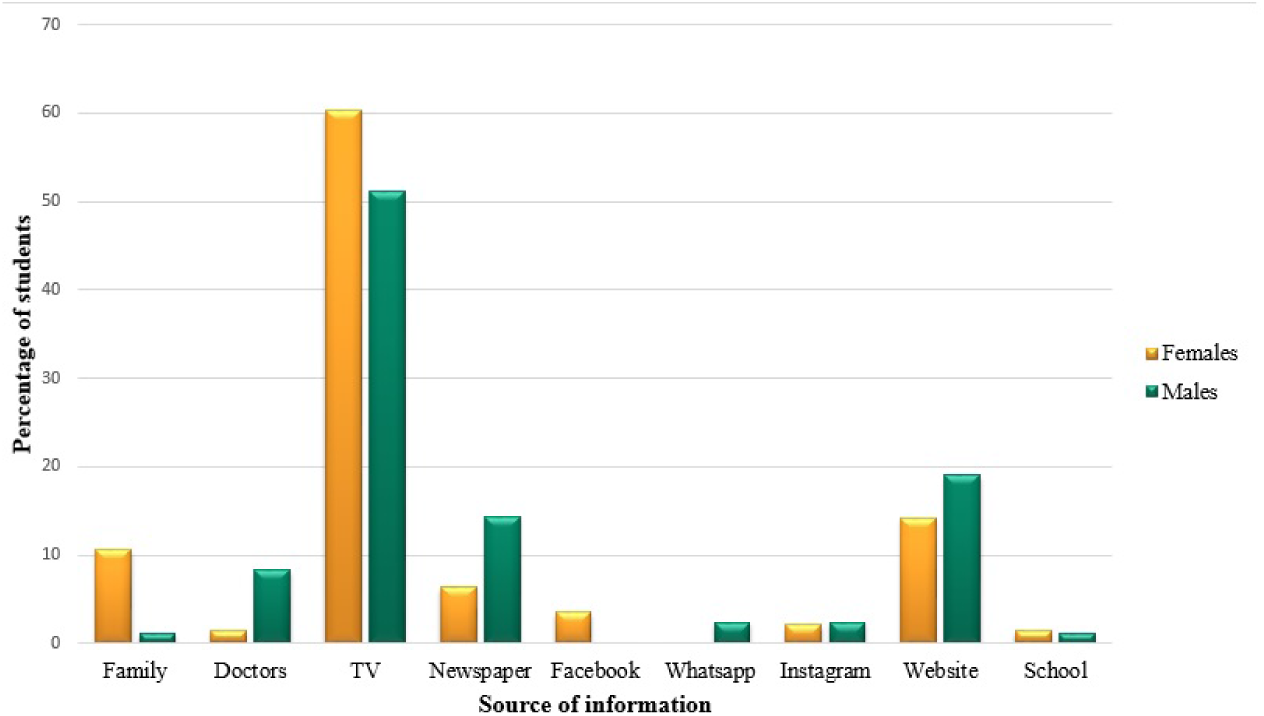
Source of knowledge about COVID-19. The source of the individuals’ information about COVID-19 includes social media and internet, news media (TV, magazines, newspapers), family, friends, school and health-care providers, such as doctors.

### 3.5 Activities regarding COVID-19

Figure 5 shows which social activity students missed most in the lockdown period. These activities are similar to two genders. The activity most affected is the possibility to visit family or friends (about 70%). In second place, students answer about the possibility to going to school. It is interesting is that the two activities more favourite by male students: going to the park and playing a sport.

**Figure 5.**
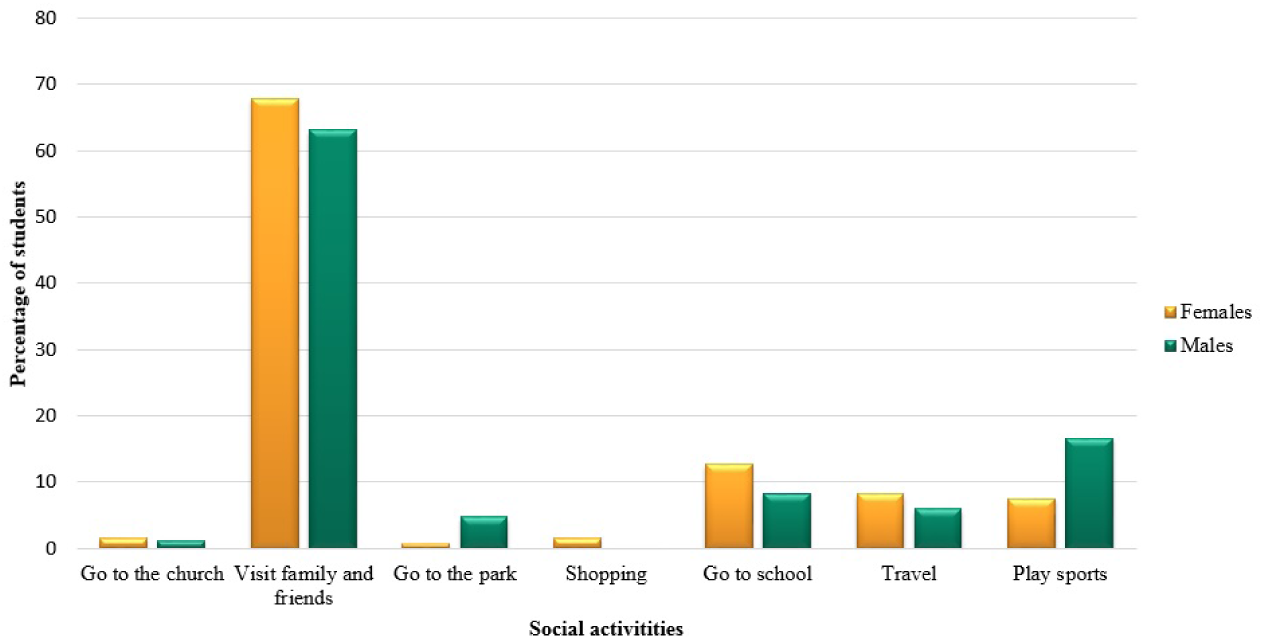
Social activities. The activity most affected is the possibility to visit family or friends (about 70%). In second place, students answer about the possibility to going to school.

Students have spent most of their free time on school online activities (see Figure 6). Italy is one of the first country in Eruope that immediately stars a new teaching method, called *DaD*. It is introduced by the Minister of Education as distance learning solution to face the impossibility of going to class at school, thus allowing the normal conduct of school lessons. It is based on remote lessons in platforms such as Zoom or Meet, upload of sources and materials on web and formal or informal communication on Whats App and Telegram groups [26]. It is interesting to note that although the students declare that they spend most of the hours online in the presence of teachers, they have previously answer that their main source of information is TV and the lesser school. An other important phenomenon is to highlight that the major of students has spent its time watching a film or chatting and video-calling. The presence of technology in these lock-down months has been a key point of their knowledge and maturation phenomenon [27]. Reading a book is one of the activities chosen in a clear minority.

**Figure 6.**
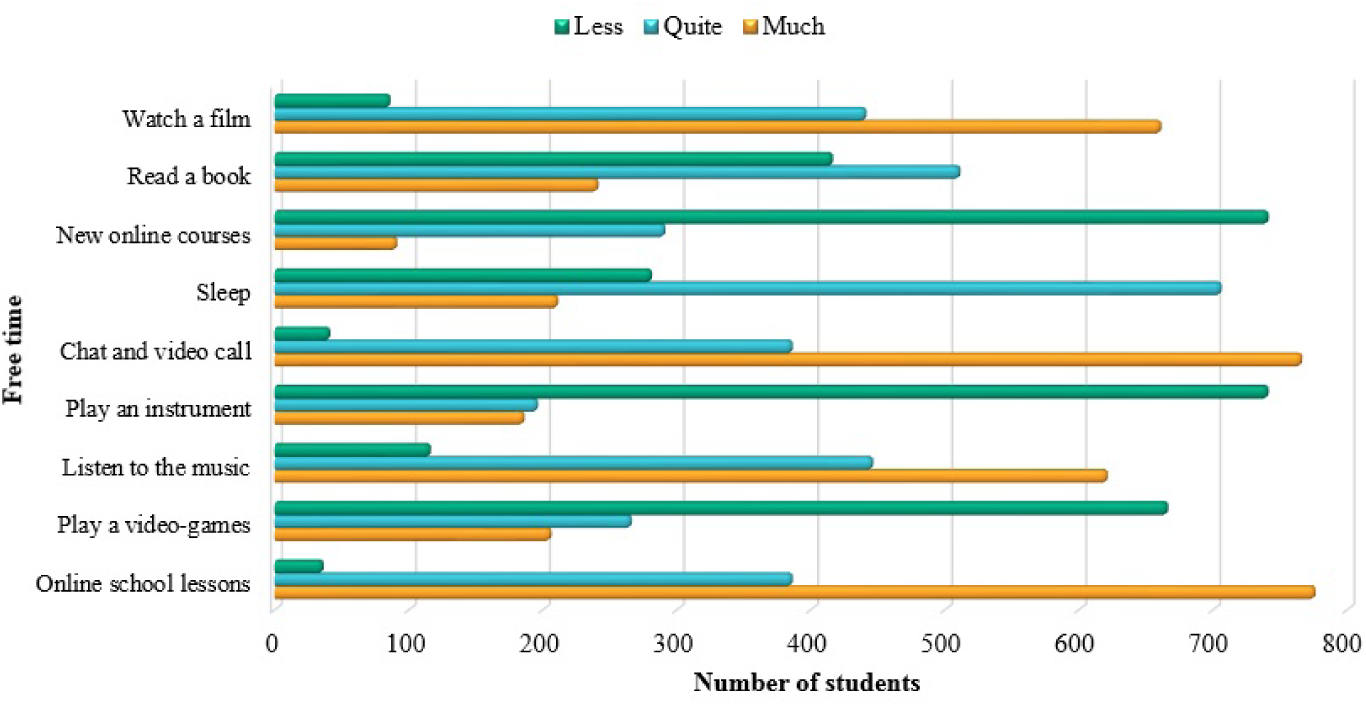
Free time. The students declare that they spend most of free time in *DaD* (online school lessons). A lot of them has spent its time watching a film or chatting and video-calling.

## Discussion

To the best of our knowledge, this is the first study in Italy investigating the KAP towards COVID-19 among the high school student population. The questionnaire regarding the knowledge of the participants about COVID-19 is divided into two sections: demographic information and KAP. In the second section we evaluate their knowledge about the characteristics of the disease and regard what they know about the mode of transmission and method of protection against virus transmission.

Out of 2380 students who answer the questionnaires, about 60% are female and the remaining male. Students attend high school from five areas of Italy: Lombardia (Milan), Lazio (Rome and Frosinone), Puglia (Lecce), Campania (Naples) and Calabria (Catanzaro). The age of participants ranges from 14 to 19 years old, with a mean age equal to 17 and SD 3.64. More than half of them attends scientific or classical high school.

The findings suggest a quite good level of perception about the disease risk. Based on our results, the majority of the general population has knowledge about the existance of virus COVID-19. The majority of students reported nasal congestion and fever as factors in the clinical presentation of COVID-19. Apart from reporting dry cough, knowledge about clinical presentation of is generally similar in the two genders. We remake that in the site of Minister of Healt they report the most common symptoms such as dry cough, flu and wearness, but not nasal congestion, that a quarter of students responds. For what concerns mode of transmission, the most frequently source reported is via coughing and sneezing.

Instead, the most frequently reported method of protection against virus transmission is correct the hand washing. Unfortunately, general information about number of COVID-19 are incorrect. We know that estimating the correct number of deaths in world and in Italy is a sensitive topic, but only 15% of students identify at least the order of measurement of these data. The importance of correct information on the numbers of this pandemic is precisely one of the key points on which to focus our attention in this research.

In the other part of the questionnaire, we ask for generic information about COVID-19 vaccine. Although no vaccine has completed clinical trials, there are multiple attempts in progress to develop such a vaccine. The major part of students answer that does not still exist a vaccine, only 5% answers that they do not know and the remaining 8.3% that vaccine exists and that soon it will be possible to do it. The other two questions correlated with vaccine are about the knowledge of antibodies. About 98% claims to know antibodies, but in the next question when they have to answer how to check for antibodies, only 65% answers with blood analysis. The remaining answers pharyngeal swab or urine analysis and aa minority of 5% answers that flu is a necessary symptome to check antibodies. These three questions with uncorrect answers are related to the above discussion. It is very important for young students to have a non-superficial knowledge of COVID-19 also from a biological and scientific point of view and not only socially.

Instead, we are interested to check which are sources of knowledge about COVID-19, including social media, internet, news media (TV/video, magazines, newspapers), family, friends, school and health-care providers, such as doctors. The most frequently reported source of knowledge is television, followed by Facebook, Whatsapp and Instagram. Unlike the media, school appears to be the last learned option of forgathering knowledge.

Finally, students declare that they have spent most of their free time on school online activities. In Italy from the first days of lock-down, the Minister of Education introduces a new learning methods for students, based on remote lessons in platforms, upload of sources and materials on web and formal or informal communication on blog or online groups. The presence of technology in these lock-down months has been a key point of their knowledge and maturation phenomenon and helps students to continue their formative learning. Perhaps more attention should be given to utilizing tecnhology and particularly social media resources, especially Facebook or Whatsapp, as techniques of promoting public health education in adolescents, in collaboration with teachers and specialists.

## Acknowledgement(s)

We warmly thank all the study participants for their voluntary participation and for providing essential information.

## Notes on contributors

DS and MD conceived and designed the experiments; MD performed the experiments; DS and MD analyzed the data; DS and MD wrote the paper. Authorship must be limited to those who have contributed substantially to the work reported.

